# Conditional blastocyst complementation of a defective Foxa2 lineage efficiently promotes generation of the whole lung

**DOI:** 10.1101/2022.10.31.514628

**Authors:** Akihiro Miura, Hemanta Sarmah, Junichi Tanaka, Youngmin Hwang, Anri Sawada, Yuko Shimamura, Yinshan Fang, Dai Shimizu, Zurab Ninish, Jake Le Suer, Nicole C. Dubois, Jennifer Davis, Shinichi Toyooka, Jun Wu, Jianwen Que, Finn J. Hawkins, Chyuan-Sheng Lin, Munemasa Mori

## Abstract

Millions suffer from incurable lung diseases, and the donor lung shortage hampers organ transplants. Identifying the crucial lineage and the program for lung organogenesis could facilitate designing whole-lung bioengineering. Using lineage-tracing mice and human iPSC-derived lung-directed differentiation, we revealed that gastrulating Foxa2 lineage contributed to both lung mesenchyme and epithelium formation. Interestingly, Foxa2 lineage-derived cells in the lung mesenchyme progressively increased and occupied more than half of the mesenchyme niche, including endothelial cells, during lung development. Foxa2 promoter-driven, conditional Fgfr2 gene depletion caused the lung agenesis phenotype in mice. Importantly, wild-type donor mouse iPSCs injected into their blastocysts rescued this phenotype by complementing the Fgfr2-defective niche in the lung epithelium and mesenchyme. Donor cell is shown to replace the entire lung epithelial and robust mesenchymal niche during early chimeric lung development, resulting in efficient complementation of the nearly entire lung niche at the late stage of lung development. These results suggest that lung complementation based on the Foxa2 lineage is a unique model for the progressive mobilization of donor cells into both epithelial and mesenchymal lung niches and provides crucial insights for designing new bioengineering strategies to generate whole lungs.

## INTRODUCTION

Tissue regeneration to treat various intractable diseases has long been challenging(Hackett et al., 2010; Kemter et al., 2020; Ott et al., 2010; Petersen et al., 2010; Wang, 2019). Organ bioengineering strategy based on recellularizing tissue-specific progenitors into the decellularized scaffolds, induced pluripotent stem cell (iPSC)-derived organoids, or 3d-bioprinters are the next-generation tissue transplant therapies(Guyette et al., 2014; Kotton and Morrisey, 2014; Petersen et al., 2010; Tian et al., 2021). Even with these techniques, however, the mammalian lung is one of the most difficult organs to replicate because of its anatomical complexity and cellular diversity. It contains hundreds of airway branches and a thin micron-sized alveolar layer of inflated and well-vascularized alveoli composed of billions of cells from more than 50 different cell types(Crapo et al., 1982; Kotton and Morrisey, 2014; Stone et al., 1992; Travaglini et al., 2020). Donor organs for lung transplantation are in short supply worldwide, but the technology does not exist to generate whole lungs composed of tissue-specific epithelial and mesenchymal cells, including endothelial cells. Lungs grow fully only through natural lung development.

During development, lung epithelial and mesenchymal precursors interact to initiate an elaborate developmental program of organogenesis that includes specification, pattern formation, progenitor cell expansion, and differentiation. The lung epithelial cells are derived from the foregut, definitive endoderm (DE) derivatives, classically labeled by Sox17 and Forkhead Box A2 (Foxa2)(Green et al., 2011; Huang et al., 2014). Multiple genetic studies using Sonic Hedgehog (Shh) Cre lineage-tracing mice have also shown that the entire Nkx2-1+ lung and tracheal epithelial primordium arises from Shh^+^ DE.

The lung mesenchyme primordium is derived from Wnt2^+^ Isl1^+^ cardiopulmonary progenitors (CPP). CPP is the derivative of Osr1^+^ Nkx6-1^+^Barx1^−^ Wnt4 ^low^ foregut lung mesoderm that arises from lateral plate mesoderm (LPM)(Han et al., 2020). While DE and LPM arise from primitive streaks (PS) during gastrulation, the exact lineage origin of LPM has been a complete mystery.

Mesendoderm is a bipotent transitional state between the PS and nascent mesoderm labeled by Mixl1, Pdgfrα, and Brachyury (T) during gastrulation that can give rise to both DE and mesoderm(Hart et al., 2002; Tada et al., 2005). Although it was speculated that mesendoderm might form LPM and DE, there have been no conclusive genetic studies on whether mesendoderm gives rise to both lung epithelium and mesenchyme. Pdgfrα is expressed in the epiblast-derived mesendoderm, the primitive endoderm (PrE), and its extra-embryonic endoderm derivatives, such as parietal and visceral endoderm, around E5.5∼E7.5. Foxa2 plays a pivotal role in alveolarization and airway goblet cell expansion(Wan et al., 2004), while there was a significant knowledge gap regarding Foxa2 lineage during lung development.

Blastocyst complementation (BC) has been proposed as a promising option for tissue-specific niche complementation(Chen et al., 1993). This unique technology has been further developed into intra- and interspecies organ generations such as kidney, pancreas, and blood vessels(Hamanaka et al., 2018; Kobayashi et al., 2010; Usui et al., 2012; Yamaguchi et al., 2017). However, the production of entire organs, including tissue-specific epithelium and mesenchyme, including endothelium, was still difficult because endothelium significantly impacts the other organs. Unfortunately, even with BC, the lungs produced were non-functional and very inefficient, and in addition, the chimeric lungs contained a substantial amount of host-derived tissue(Kitahara et al., 2020; Li et al., 2021; Wen et al., 2021). Previously, we established the conditional blastocyst complementation (CBC) approach, which targets specific lineages complemented by donor pluripotent stem cells(Mori et al., 2019). Using lineage-specific drivers of lung endoderm in CBCs avoids the effects of genetic manipulation in non-target organs for the generation of empty organ niches that lead to functional chimeric lung generation(Mori et al., 2019). However, most of the lung mesenchyme and endothelium were still derived mainly from the host cells, which was the severe limitation of CBC(Mori et al., 2019). Given that the CBC approach targeted endodermal lungs, we speculated that this limitation was due to a significant gap in our knowledge of the origin of all lung cell types, especially pulmonary mesenchyme, including endothelium. In particular, the complementation of endothelium is a critical issue for overcoming hyperacute rejection after lung transplantation. To overcome this critical issue, we explored the origin and the program of whole lung epithelium and mesenchyme, the major components of the lung.

We hypothesized that targeting a single bona fide lung generative lineage (BFL) may facilitate the designing of the entire lung generation. BFL is based on the strategy for vacating the desired niche by lineage-specific gene depletion to be complemented by donor cells after injecting them into the host blastocysts. The lineage is not necessarily lung-specific but rather based on emptying or mitotically defective in the desired lineage by gene depletion.

We found that the Foxa2 lineage functions as a BFL in CBC. Although the Foxa2 lineage did not label the entire lung cells during lung development, the mitotically-defective Foxa2 lineage promoted efficient whole-lung complementation by donor cells.

## RESULTS

### Pdgfr*α*^+^ lineage during gastrulation gives rise to the entire lung mesenchyme

To determine the origin of LPM and pulmonary endothelium for the whole lung generation via BFL, we performed lung mesenchyme precursor lineage-tracing analysis using *Pdgfrα*^*CreERT2/+*^; *Rosa*^*tdTomato/+*^ mice. Surprisingly, tamoxifen injection at E5.5 labeled the entire lung mesenchyme with tdTomato at E14.5 (Figures 1A and 1B). This result suggested that the origin of the whole lung mesenchyme is the Pdgfrα lineage around early-to-mid-streak-stage embryos. tdTomato labeled the entire pulmonary mesenchyme, including Sma^+^ airway smooth muscle cells, Pdgfrβ^+^ pulmonary mesenchyme, and VE-cadherin^+^ vascular endothelial cells (Figure 1B). In addition, the Pdgfrα lineage only partially labels the lung epithelium (Figure 1B, arrows), suggesting that the contribution of the Pdgfrα lineage to the lung endoderm is low. It suggests that Pdgfrα is difficult to define as a BFL because of its low contribution to the lung epithelium.

**Figure 1.**
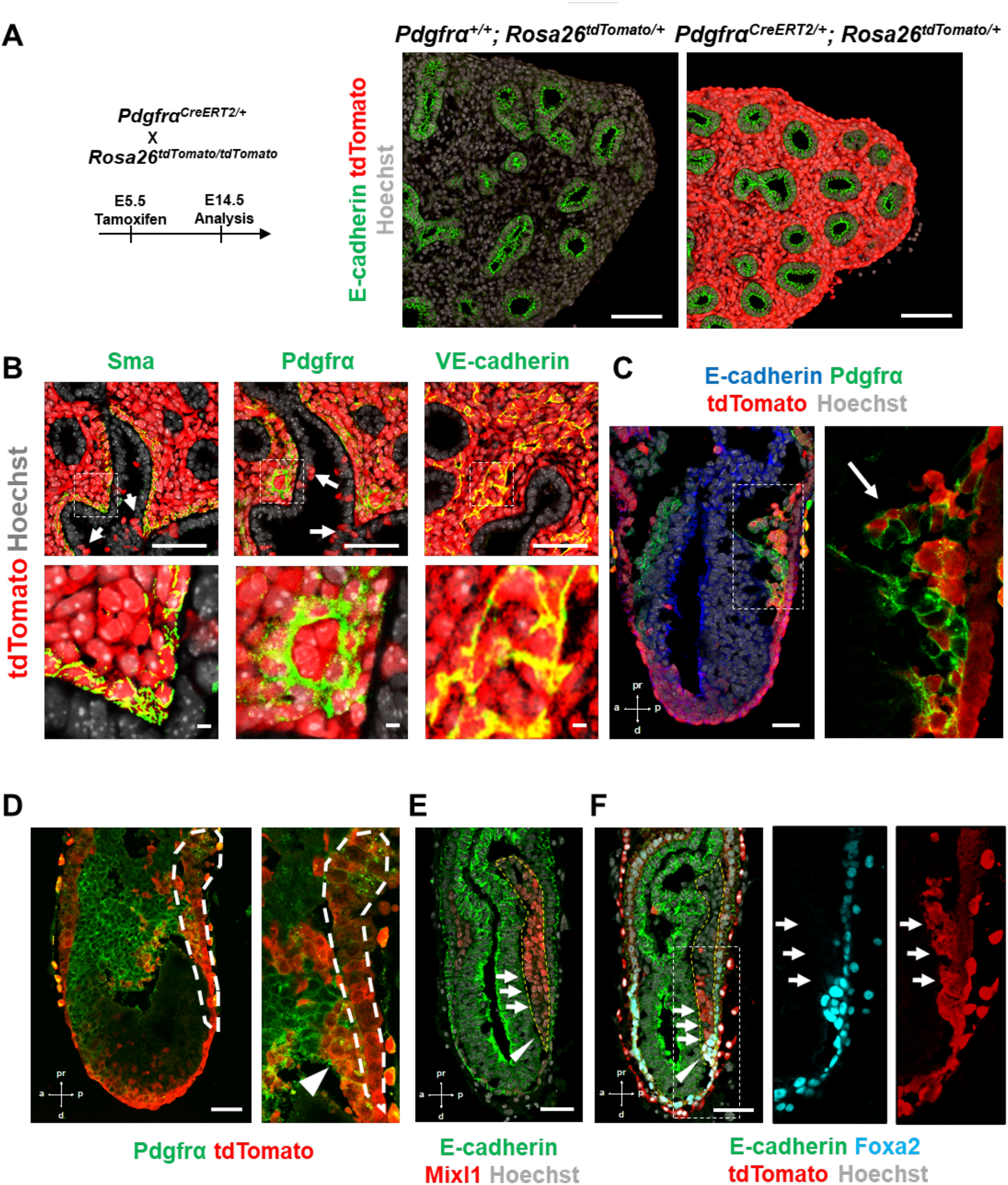
Pdgfr*α* lineage during gastrulation is the origin of the entire pulmonary mesenchyme. (A) Left: Schematic of Tamoxifen administration (A-B). Representative Immunofluorescence (IF)-confocal imaging of E14.5 *Pdgfr*α^*CreERT2/+*^; *Rosa26*^*tdTomato/+*^ lineage tracing mouse lungs (A, B). Tamoxifen administration at E5.5 labels Pdgfrα-lineage-driven tdTomato (red) in the entire lung mesenchyme (A), including Sma^+^ airway smooth muscle cells, Pdgfrα^+^ mesenchyme, and VE-cadherin^+^ capillaries (B). Pdgfrα-lineage also labeled a low proportion of epithelial cells (B, white arrows). Enlarged box: dotted box. Cre^−^ littermate control (A, middle panel) (*n = 3* per group). Scale bars: A, B = 100, 500μm. (C-F) Representative IF-confocal imaging of E6.25 (C), E7.0 (D), and E6.5 (E, F) from *Foxa2*^*Cre/+*^; *Rosa26*^*tdTomato/+*^ mice. E-cadherin indicates epiblasts (C-F): (C) Foxa2-lineage (red) labeled Pdgfrα (green) expressing mesendoderm (arrow). (D) Foxa2-lineage labeled Pdgfrα^+^ cells and ingresses from primitive streak (PS) (dotted lines) to nascent mesoderm regions (arrowhead). (E) Mixl1 (red) expression in PS (yellow dotted area). (F) Sequential section of E: Enlarged box: Foxa2-lineage (red) marked the distal portion of the arteriolarizing PS that is a part of the MIxl1^dim+^ mesendoderm (E-F, arrowheads). (*n = 3* per group). Scale bars = 50μm.

### Foxa2 lineage labels Pdgfr*α*^+^ mesendoderm niche during mouse development

Single-cell RNA-seq (scRNA-seq) analysis using Foxa2-Venus fusion protein reporter mice indicated that the Foxa2 lineage might give rise to LPM and DE (Scheibner et al., 2021). Given that Foxa2 and Pdgfrα are expressed during the conversion from mesendoderm to mesenchyme(Artus et al., 2010; Kopper and Benvenisty, 2012; Scheibner et al., 2021; Tada et al., 2005), we used Foxa2-lineage tracing mice (*Foxa2*^*Cre/+*^; *Rosa*^*tdTomato/+*^)(Horn et al., 2012) to determine whether the Foxa2-lineage would label Mixl1^+^ or Pdgfrα^+^ mesendoderm around E6.25∼7.0 during gastrulation. Notably, Foxa2-lineage-driven tdTomato labels a part of the Pdgfrα^+^ mesendoderm of the primitive streak (PS) around E6.25∼E6.5 relatively early-streak-stage embryos (Figure 1C, arrow). In the E6.5∼E7.0 mid-to-late-streak-stage PS region (Figure 1D, dotted lines), Foxa2-lineage-derived tdTomato were found in the PS and Pdgfrα^+^ adjacent nascent mesoderm, suggesting that Foxa2 lineage-labeled Pdgfrα^+^ cells ingresses from PS to nascent mesoderm regions (Figure 1D, arrowheads). These data are reminiscent of the definition of mesendoderm (Hart et al., 2002; Tada et al., 2005). Our result indicates that the Foxa2 lineage-labeled Pdgfrα^+^ cells are the mesendoderm, most likely the derivative of posterior epiblasts, since Foxa2 is expressed only in the posterior epiblasts before gastrulation (Scheibner et al., 2021). Further immunostaining analysis of the sequential sections of mid-streak-stage E6.5∼6.75 embryos confirmed that the Foxa2 lineage appeared in the distal portion of Mixl1-weakly-positive mesendoderm (Figures 1E and 1F, arrows) at anterior primitive streak (APS) besides the Foxa2 protein-expressing DE (Figures 1E and 1F, arrowheads). Based on our Pdgfrα^+^ lineage tracing data, we further examined whether Foxa2-lineage would label lung mesenchyme.

### Foxa2-lineage labeling increased during lung development, leading to occupy the entire lung epithelium and half of the lung mesenchyme, including lung endothelium

Foxa2-lineage tracing mice (*Foxa2*^*Cre/+*^; *Rosa*^*tdTomato/+*^) faithfully target Nkx2-5^+^ cardiac progenitors associated with the origin of Wnt2^+^ Isl1^+^ CPP(Bardot et al., 2017; Peng et al., 2013) lung mesenchyme. However, there was no conclusive evidence of whether Foxa2-lineage can give rise to Wnt2^+^ Isl1^+^ CPP(Bardot et al., 2017; Peng et al., 2013). Using the Foxa2-lineage tracing mice, we found from immunostaining that Foxa2 lineage labeling occupies the entire lung epithelium and most of the lung mesenchyme at E16.5 (Figures 2A-C). Quantitative analyses by flow cytometry in the E14.5 developing lungs of Foxa2-lineage tracing mice showed that Foxa2-lineage labeled almost the entire lung epithelium (93.3%± 2.00) and the partial lung mesenchyme (24.2%± 5.51), including endothelial cells (16.8 %± 5.12) (Figure 2D). Contrary to expectations, Foxa2-lineage labeled cells increased dramatically throughout lung development (Figure 2E). In adulthood, the Foxa2-lineage labeling reached about 99.97%± 0.0578 in lung epithelium, 51.27%± 1.72 in lung mesenchyme, and 59.97%± 8.44 in lung vascular endothelial cells, with more than two-fold change in the lung endothelium, compared with E14.5 (Figure 2E). Morphometric analysis of immunostaining further confirmed that Foxa2 lineage marked about 30% of E14.5 multiple cell types of lung mesenchyme: Sma^+^ smooth muscle cells (24.9%± 8.50), VE-cadherin^+^ (39.3%± 12.5) or Pecam1^+^ (36.6%± 11.1) endothelial cells, and Pdgfrβ^+^ pericytes of the pulmonary arteries and pulmonary veins (41.4%± 10.5) (Figures 1F and 1G). Interestingly, no clear Foxa2 protein level expression was observed in the mesenchyme of the embryonic lungs (Figure S1A), consistent with previously reported Foxa2 protein expression patterns(Wan et al., 2004). However, the LungMAP deposited single-cell RNA-seq database analysis showed a sporadic Foxa2 transcriptional expression pattern in developing lung mesenchyme, particularly in proliferating endothelium, on E15.5 and E17.5 (Figure S1B). We confirmed *Foxa2* transcriptional expression in lung mesenchyme by in situ hybridization analysis on E18.5 (Figure S1C and S1D). The expression was detected mostly in tdTomato positive but low frequency in the negative cells, which indicates that lung mesenchyme spontaneously expressed *Foxa2* at a transcriptional level and turn on tdTomato during lung development. We also sorted the cell fraction of CD45^−^ CD31^−^ EPCAM^−^ tdTomato^+^ and CD45^−^ CD31^−^ EPCAM^−^ tdTomato^−^ from developing lung mesenchyme at E18.5. We observed a slight increase in the relative expression of *Foxa2* in the tdTomato^+^ fraction of embryonic lung mesenchyme, which most likely contributed to the labeling in the Foxa2-lineage tracing mice (Figure S1E). These results suggest pulmonary mesenchymal progenitor cells turned on Foxa2 expression slightly at the mRNA level rather than the protein level, which led to a gradual increase in Foxa2 lineage labeling. Furthermore, tdTomato^+^ endothelial cells were found to have a slightly higher proliferative capacity than tdTomato^−^ cells (Figure S1F). These results suggest that Foxa2 lineage^+^ pulmonary mesenchymal appeared during lung development, and lineage labeling gradually increased compared to Foxa2^−^ lung mesenchyme progenitors throughout lung development.

**Figure 2.**
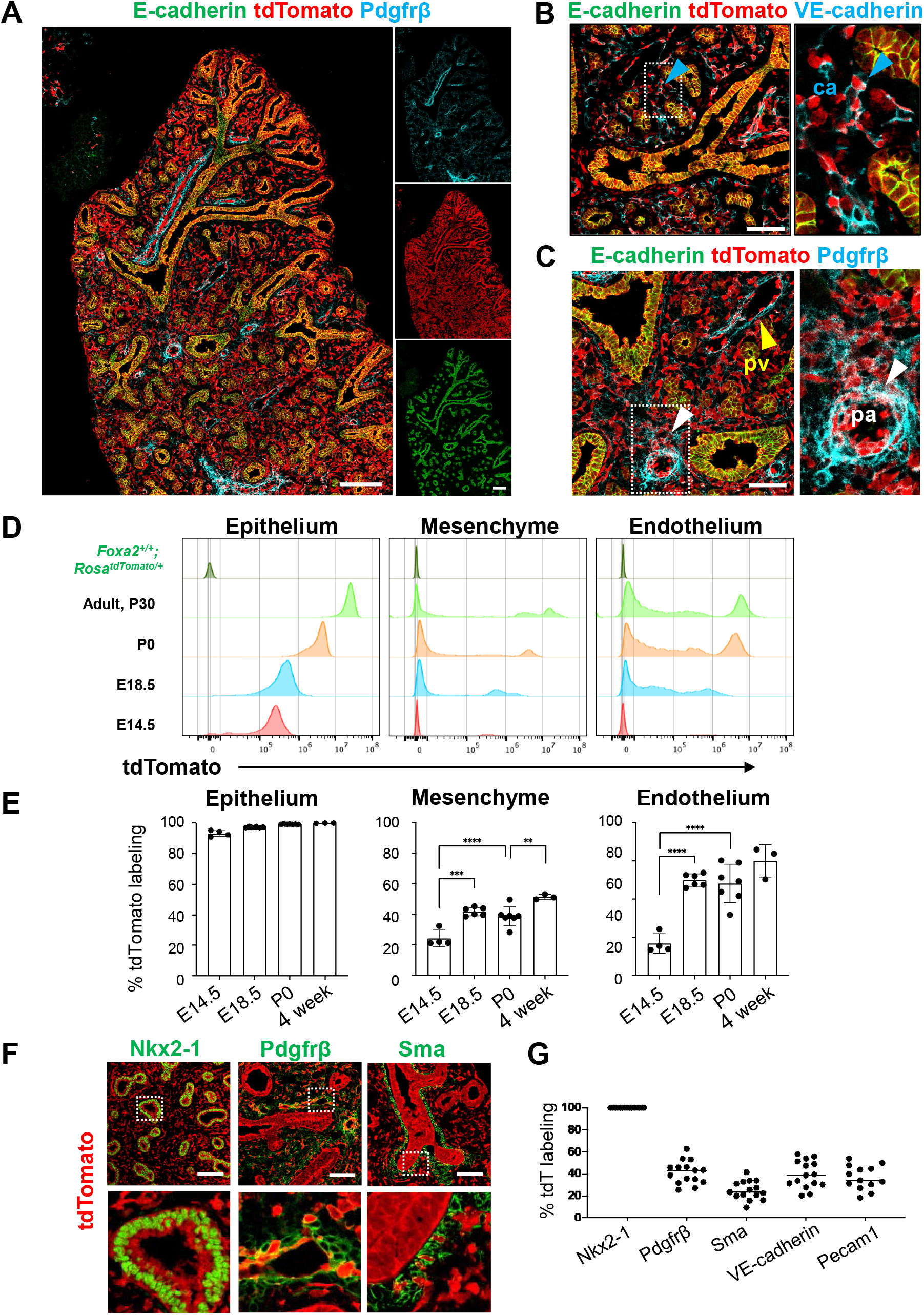
Foxa2-lineage gradually increased during lung development and labeled the entire lung epithelium and half of the mesenchyme. (A-C) IF-confocal imaging of E16.5 *Foxa2*^*Cre/+*^; *Rosa*^*tdTomato/+*^ embryonic lung: (A) Foxa2-lineage (red) labeled E-cadherin^+^ lung epithelium (green) entirely and Pfgfrβ^+^ mesenchyme (cyan) partially. (B) Foxa2-lineage partially labeled VE-cadherin^+^ capillary (ca) (enlarged box, blue arrowhead). (C) Foxa2-lineage labeled Pdgfrβ^+^ smooth muscle cells of the pulmonary artery (pa) (enlarged box, white arrowhead) and pulmonary vein (pv, yellow arrowhead). Scale bars (A), (B), and (C) = 200μm, 100μm, and 100μm, respectively. (D and E) Representative histograms and the graphs of FCM quantitative analyses for CD31^−^ Epcam^+^ lung epithelium, CD31^−^Epcam^−^ mesenchyme, and CD31^+^Epcam^−^ endothelium at E14.5, E18.5, P0, and four weeks adult (n = 4, 6, 7 and 3, independent biological replicates, respectively) of *Foxa2*^*Cre/+*^; *Rosa*^*tdTomato/+*^ mouse lungs. The gradual increase of % tdTomato^+^ lineage labeling in both lung mesenchyme and endothelium. Statistical analysis: one-way ANOVA with the Tukey post hoc test.; statistically significant if \**P* < 0.05, \*\**P* < 0.01, \*\*\**P* < 0.001, \*\*\*\**P* < 0.0001. (F-G) IF-confocal imaging of E14.5 *Foxa2*^*Cre/+*^; *Rosa*^*tdTomato/+*^ embryonic lungs. tdTomato labeled entirely with lung epithelial markers Nkx2-1(left) but a relatively low proportion of mesenchyme (Pdgfrβ: middle, and Sma: right). (n = 3 per group). Scale bars = 50μm. Graphs in (G): The morphometric analysis: % of Foxa2-lineage labeling in Nkx2-1^+^epithelial, Pdgfrβ^+^mesenchyme, Sma^+^airway smooth muscle, VE-Cadherin^+^ capillaries, or PECAM1^+^ arteries from E14.5 *Foxa2*^*Cre/+*^; *Rosa*^*tdTomato/+*^ lungs. (n = 3 per biological replicates, 5 fields per group).

### Co-development of endodermal and mesodermal lung progenitors derived from MXIL1^+^ PDGFR*α*^+^ FOXA2^+^ mesendoderm in the directed differentiation protocol using hiPSC

To determine whether Foxa2 or Pdgfrα mesoderm is an evolutionarily well-conserved niche that can give rise to both pulmonary endoderm and mesoderm, we modified a previously reported protocol to establish a pulmonary endoderm-mesoderm co-developmentally directed differentiation protocol(Chen et al., 2017; Gotoh et al., 2014; Hawkins et al., 2017; Huang et al., 2014; Konishi et al., 2016) (Figure 3A). With this optimized protocol, various hiPSC lines were found to efficiently induce a lung bud-like appearance, indicated by NKX2-1+, in lung epithelial cells(Gotoh et al., 2014; Konishi et al., 2016). We found that on day 10, TBX4^+^ lung mesenchyme emerged and surrounded the NKX2-1^+^SOX9^+^ lung epithelium (Figure 3B and 3C). The qPCR kinetics analyses across the time point further supported the appearance of lung mesenchyme, represented by LPM marker expression peaked on day6∼8; *OSR1, FGF10, BMP4, PDGFRα*, and smooth muscle cell markers peaked on day8∼10; *ACTA2* and *PDGFRβ*, and CPP markers peaked on day8∼12; *ISL1, WNT2, FOXF1*, and *TBX4*, peaked on days 10∼14 simultaneously with pulmonary epithelial markers, *NKX2-1* and *CPM* (Figure 3D).

**Figure 3.**
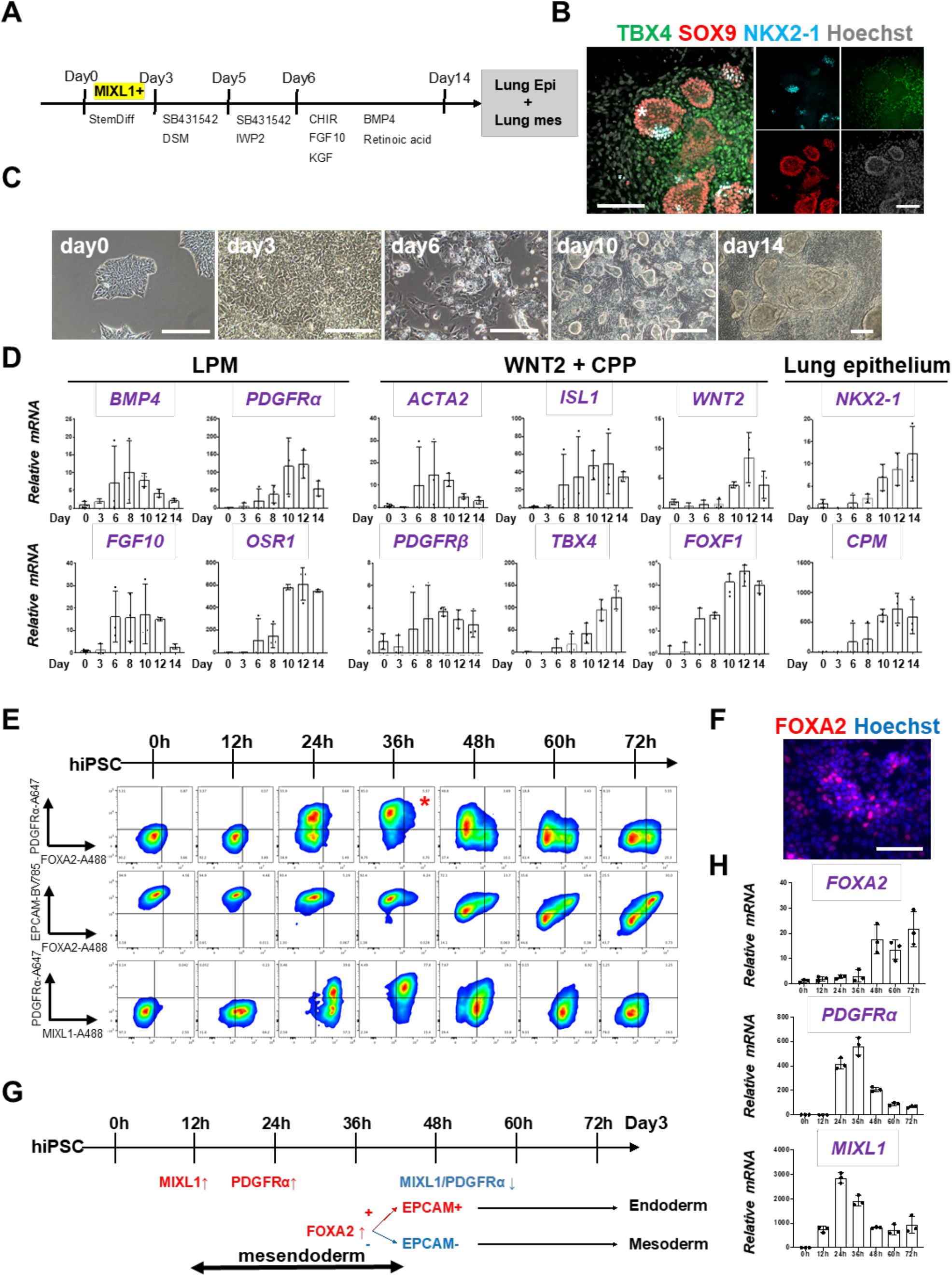
Co-development of endodermal and mesodermal lung progenitors derived from MXIL1^+^ PDGFR*α*^+^ FOXA2^+^ mesendoderm in the directed differentiation protocol using hiPSC. (A) Schematic culture protocol of hiPSC-derived endodermal and mesodermal lung progenitor cell co-differentiation. (B) Representative IF-confocal imaging of differentiating hiPSCs at day 10 culture. Lung epithelium (NKX2-1), distal lung bud epithelium (SOX9), mesenchyme (TBX4), and nucleus (Hoechst) markers. The budding structures expressed SOX9 and partially NKX2-1 (asterisk), and monolayer cells expressed TBX4. (C) Representative phase-contrast images of the directed differentiation time course (D) qRT-PCR analyses of lung mesenchyme and epithelium markers in time course according to the protocol shown in (A). Each plot showed a different biological experiment (n = 3 independent experiments). (E) FCM-based protein kinetic analyses during DE and LPM induction; MIXL1 expression preceded compared to PDGFRα or FOXA2. FOXA2 appearance in the subset of the PDGFRα^+^ population (red asterisk). (*n = 3* independent experiments) (F) Representative IF imaging of 36 hours-cultured hiPSCs. (G) Schematic summary of flow cytometry analysis. (H) qRT-PCR analyses further confirmed the preceded *MIXL1* induction and subsequent expression of *PDGFRα* and *FOXA2*. All graphs: Data normalized by undifferentiated iPSCs. Each plot showed a different biological experiment (n = 3 independent experiments).. Error bars represent mean ±SD.

In this differentiation protocol, NKX2-1^+^ lung endoderm and WNT2+TBX4^+^ lung mesoderm were derived from the anteroventral endoderm and mesoderm at day 15 after Activin-mediated definitive endoderm and LPM induction, respectively(Chen et al., 2017; Huang et al., 2014). During primordial streak induction from day0 to day3, cell surface markers of PDGFRα and EPCAM and intracellular FOXA2 and MIXL1 kinetics were analyzed by flow cytometry every 12 hours (Figure 3E). Briefly, 12 hours after the Activin induction, more than 60% of the EPCAM^+^PDGFRα^−^ primitive streak first turned on MIXL1, the mesendoderm marker(Hart et al., 2002; Tada et al., 2005). Subsequently, the epithelial-mesenchymal transition occurred 24 hours later, as represented by the PDGFRα induction in EPCAM^+^MIXL1^+^ mesendoderm. After 36 hours, more than 90% of MIXL1+EPCAM+ mesoderm cells expressed PDGFRα. At the same time, expression of FOXA2 appeared in some of those mesoderm cells (Figures 3E and 3F). Thereafter, PDGFRα expression decreased, and 72 hours later, mutual FOXA2 induction appeared when EPCAM^+^FOXA2^+^ DE and EPCAM^−^FOXA2^−^ LPM were presented (Figure 3G). The dynamics of MIXL1, PDGFRα, and FOXA2 were further revealed by qPCR analysis (Figure 3H). These results suggest that PDGFRα^+^ and FOXA2^+^ lineage are redundant but distinct stages of mesoderm development. It can give rise to both endoderm and mesoderm lung cells conserved in mouse development and human-directed differentiation (Figure S4A).

### Foxa2-driven Fgfr2 conditional knockout showed a lung agenesis phenotype

The evolutionary-conserved Foxa2-lineage^+^ mesendoderm forms endodermal and mesodermal lung niches, suggesting that it may function as a BFL to generate the entire lungs after donor cells are injected into the vacant niche in the Foxa2 lineage. To explore this possibility, CBC was performed using Foxa2-driven Fgfr2-conditional knockout mice (*Foxa2*^*Cre/+*^; *Fgfr2*^*flox/flox*^, hereafter, *Foxa2*^*Cre/+*^; *Fgfr2*^*cnull*^). Mitotic signaling via Fgfr2 is required for both lung epithelium and mesenchyme, and systemic knockout mice of Fgf10 or Fgfr2 exhibit a phenotype of lung agenesis(Arman et al., 1999; De Langhe et al., 2006; De Moerlooze et al., 2000; Sekine et al., 1999). Based on the results of Foxa2 lineage tracking and the need for Fgfr2 signaling, it was predicted that *Foxa2*^*Cre/+*^; *Fgfr2*^*cnull*^ mice would be used to generate vacant niches in both lung epithelium and mesenchyme. Indeed, they exhibited a lung agenesis phenotype (Figures S2A-S2C). However, we did not observe agenesis phenotype in other major internal organs related to the Fgfr2 systemic knockout phenotype (Figure S2D) (Arman et al., 1999; De Langhe et al., 2006; De Moerlooze et al., 2000; Sekine et al., 1999).

### Generation of the entire lungs in *Foxa2*-driven *Fgfr2*-deficient mice via CBC

To examine whether donor cells complement the lung agenesis phenotype, we generated nGFP^+^iPSCs from Rosa^nT-nG^ mice (hereafter, nGFP^+^iPSCs) via Sendai virus-mediated reprogramming(Huang et al., 2014). nGFP^+^iPSCs were injected into mouse blastocysts (Figure 4A), and chimerism was analyzed at E17.5. Strikingly, donor nGFP^+^iPSCs generated whole lungs in *Foxa2*^*Cre/+*^; *Fgfr2*^*cnull*^ mice, but general chimerism in other organs were diverse (Figure 4B, S3A, and Table 1). Importantly, almost the entire lung epithelial, mesenchymal, and endothelial cell population at E17.5 was composed exclusively of nGFP^+^iPSCs (Figures 4C and S3B). In contrast, wild-type, Shh-driven heterozygous, or knockout mice showed about 50-70% chimerism in the mesenchymal and endothelial lineages of the lung (Figures 4C and 4D). Interestingly, although tdTomato^+^ Fgfr2 knockout mesenchymal cells remained in early lung development at E14.5 (Figure S3C and Table 2), the percentage of Ki67^+^ proliferating cells was significantly higher in GFP^+^ donor cells compared to tdTomato^+^ host cells (Figures 4E – 4G). These results suggest that generating an Fgfr2 deficient niche in the host Foxa2 lineage-derived lung mesenchyme is efficient for donor iPSCs recruitment in the defective Foxa2 lineage mesenchyme niches.

**Table 1.**
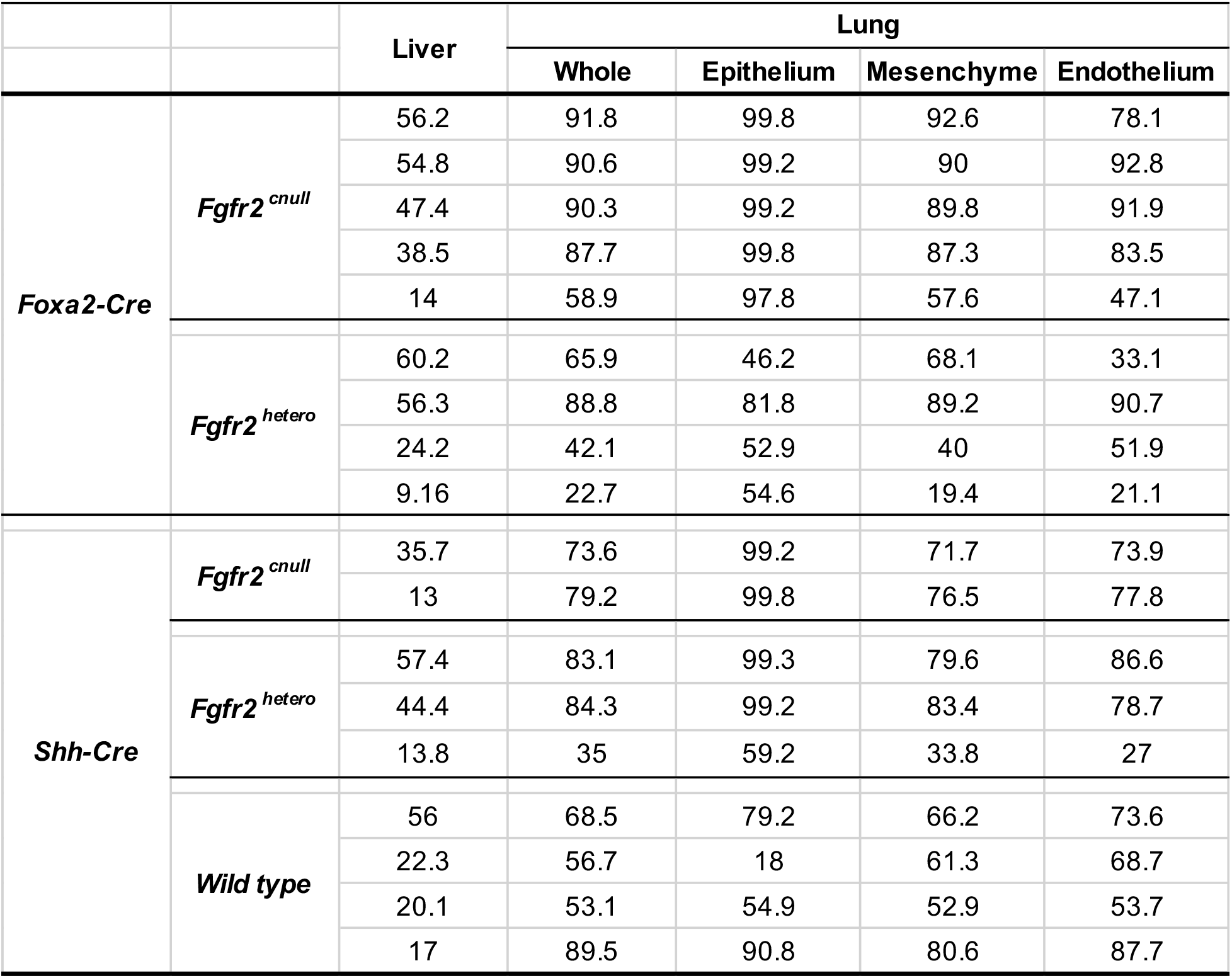
E17.5 Chimerism of Foxa2 promoter-driven conditional blastocyst complementation.

**Table 2.**
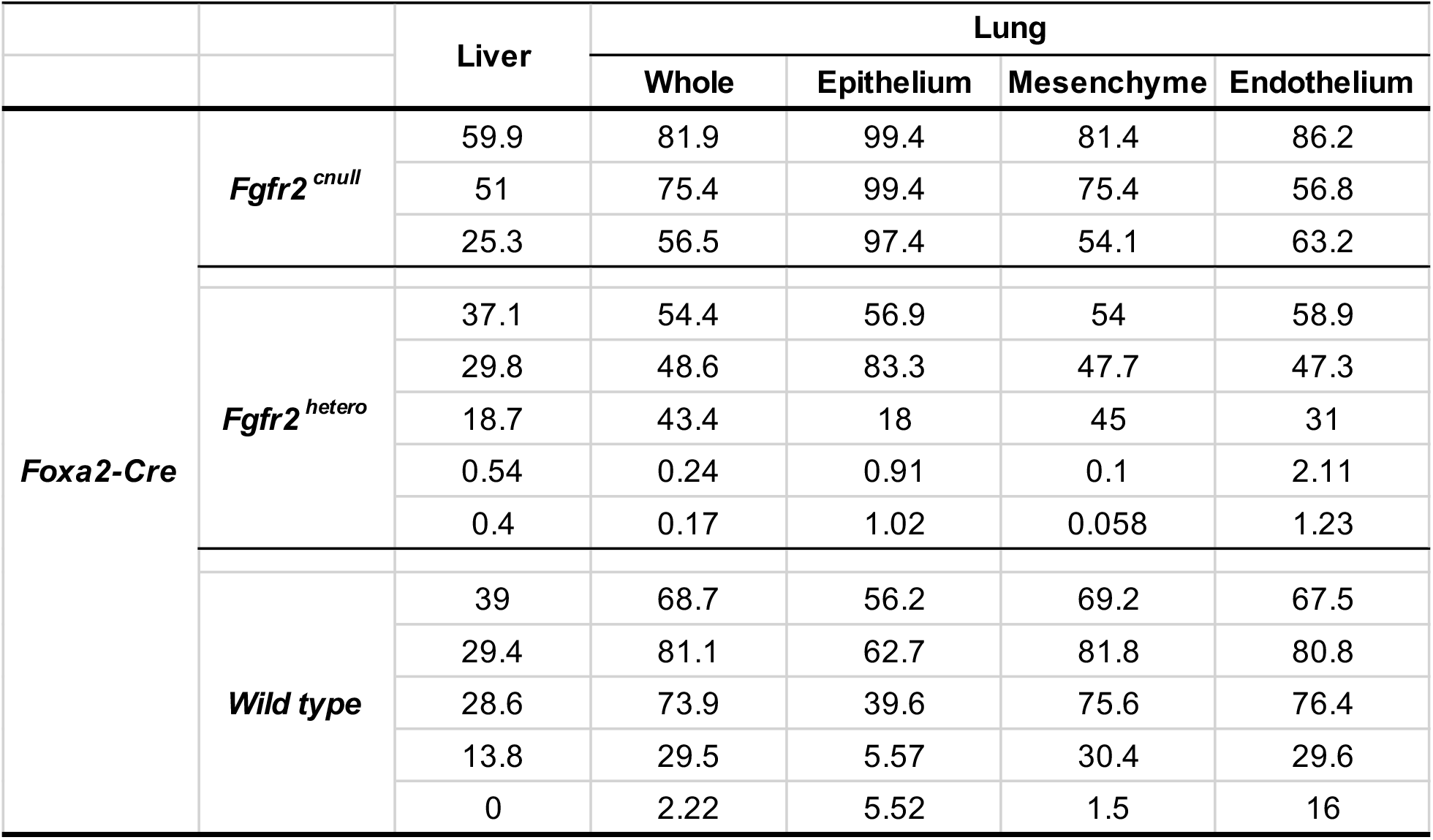
E14.5 Chimerism of Foxa2 promoter-driven conditional blastocyst complementation.

**Figure 4.**
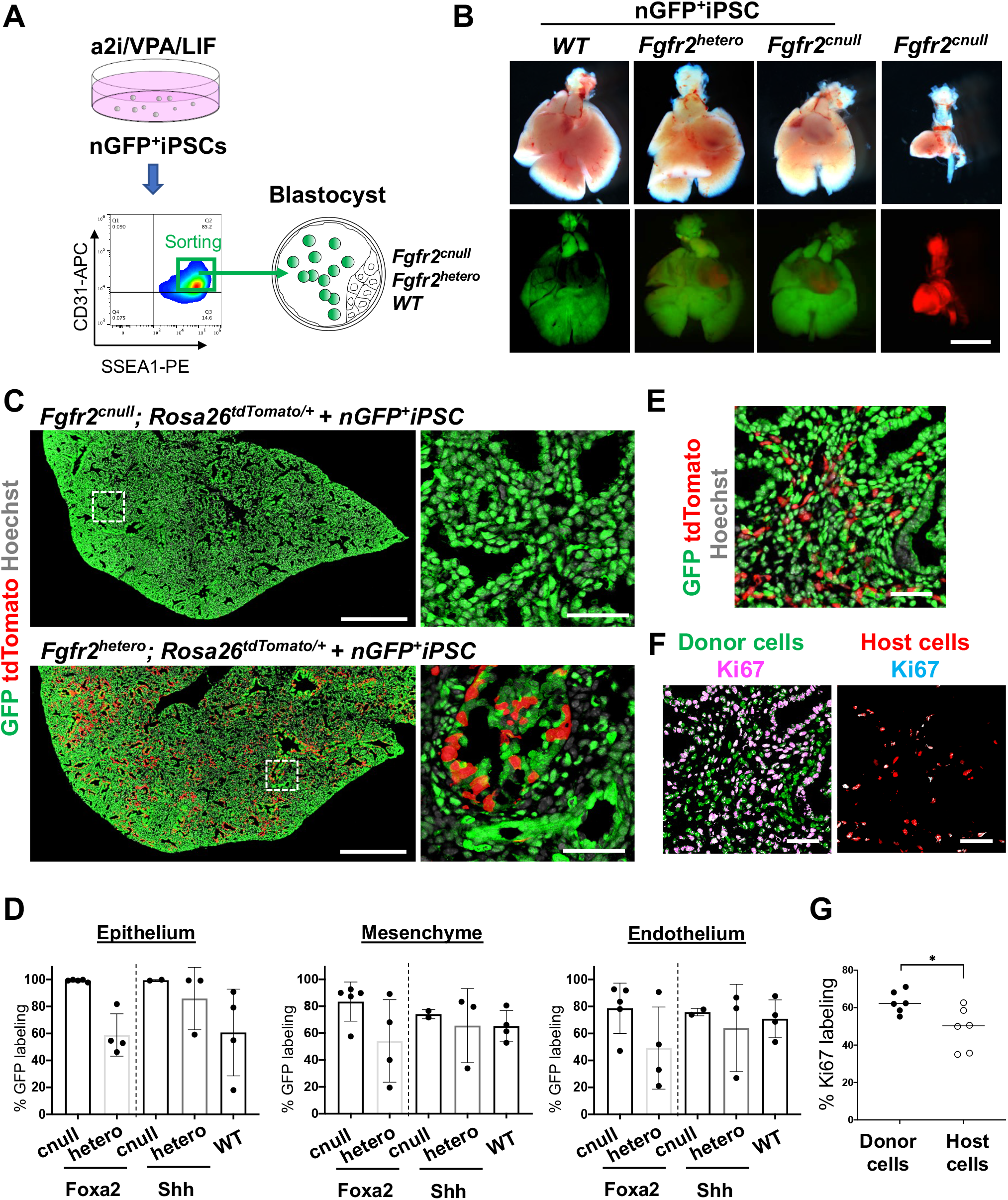
Generation of the entire lungs in *Foxa2*-driven *Fgfr2*-deficient mice via CBC. (A) Schema of CBC experiment: a2i/VPA/LIF-treated SSEA1^high^ CD31^high^ nGFP^+^iPSCs were sorted and injected into WT, *Fgfr2*^*hetero*^ (heterozygous: *Foxa2*^*cre/+*^; *Fgfr2*^*flox/+*^; *Rosa26*^*tdTomato/+*^), and *Fgfr2*^*cnull*^ (homozygous: *Foxa2*^*cre/+*^; *Fgfr2*^*flox/flox*^; *Rosa26*^*tdTomato/+*^) blastocysts. (B) Gross morphology, GFP (green: donor nGFP^+^iPSCs-derived signals), and tdTomato (host Foxa2-lineage-derived signals) fluorescence of freshly isolated lungs from E17.5 chimeric *WT*(left), *Fgfr2*^*hetero*^ (left middle), and *Fgfr2*^*cnull*^ (right middle) that were injected with nGFP^+^iPSCs. Control: littermate *Fgfr2*^*cnull*^ mouse without nGFP^+^iPSCs injection (right). (C) IF-confocal imaging of E17.5 *Fgfr2*^*cnull*^ or *Fgfr2*^*hetero*^ lungs injected with nGFP^+^iPSCs. Dotted lines: enlarged images: Compared with littermate control holding host-derived cells (red), E17.5 *Fgfr2*^*cnull*^ lungs were entirely composed of donor-derived nGFP^+^ cells (green). (D) Graphs: % GFP in CD31^−^Epcam^+^ lung epithelium, CD31^−^Epcam^−^ mesenchyme, and CD31^+^Epcam^−^ endothelium analyzed by flow cytometry. Each plot: a different biological animal. *Foxa2*^*Cre/+*^; *Fgfr2*^*cnull*^; *Rosa*^*tdTomato*^ (*n=5*, independent biological replicates), *Foxa2*^*Cre/+*^; *Fgfr2*^*hetero*^; *Rosa*^*tdTomato/+*^(*n=4*), *Shh*^*Cre/+*^; *Fgfr2*^*cnull*^; *Rosa*^*tdTomato/+*^ (*n=2*), *Shh*^*Cre/+*^; *Fgfr2*^*hetero*^; *Rosa*^*tdTomato/+*^(*n=3*), and *WT* (*n=4*). (E and F) Representative IF staining of E14.5 lung of *Fgfr2*^*cnull*^. GFP and tdTomato indicate donor and host-derived cells, respectively. (E- F) Residual tdTomato^+^ host cells in E14.5 *Foxa2*^*Cre/+*^; *Fgfr2*^*cnull*^ chimeric lungs. (F) Split images of visualizing GFP^+^ donor cells and tdTomato^+^ host cells, co-stained with Ki67. (G) Graphs: % Ki67 labeling in mesenchymal cells of E14.5 *Foxa2*^*Cre/+*^; *Fgfr2*^*cnull*^ chimeric lungs. Statistical analyses: paired Student’s t-test, significance at \**P* < 0.05, ns: non-significant. Scale bars: B, C (left, right), E, F =1mm, 500μm, 50μm, 20μm, 20μm.

## DISCUSSION

The presence of committed organ precursors capable of contributing to multi-embryonic layers after pluripotent epiblast formation has been assumed, but the identity and origin of the lung precursors needed to be better defined. Although the Foxa2 lineage could not give rise to the entire lung cells that served as a BFL, we have identified a Foxa2 lineage that provides a significant stepping stone for facilitating whole lung generation during mouse development. Since donor cells formed about 50∼80% chimerism in the complemented lung mesenchyme niches at E14.5 (Table 1) and endogenous Foxa2 lineage forms only 20%∼30% in the lung mesenchyme at E14.5, Fgfr2 defects at the Foxa2 lineage in early lung development are important but not sufficient for the complete lung mesenchyme niche complementation. We showed that losing Fgfr2 expression in Foxa2 lineage cells resulted in lower proliferative ability than donor cells during the lung complementation (Figure 4F).

Lung epithelial cell precursors were well-known to be Shh^+^ DE in the lung development field(Cardoso and Kotton, 2008; Christodoulou et al., 2011; Harris et al., 2006; Kadzik and Morrisey, 2012; Tian et al., 2011; Weaver et al., 1999; Xing et al., 2008), and indeed, the Shh-lineage traces putative DE-derived epithelial lineage but little lung mesenchyme (Figure S3D). Targeting the endodermal lung lineage driven by Shh was sufficient for lung epithelial complementation but insufficient to generate whole lungs, and host-derived cells remained substantially in the mesodermal lung component (Mori et al., 2019).

In contrast to lung epithelium precursors, the orderly commitment and the origin of the entire pulmonary mesenchyme were not defined well. We showed that gastrulating Pdgfrα-lineage is the origin of the entire lung mesoderm, including endothelium (Figures 1A and 1B)(Mori et al., 2019). Furthermore, Foxa2 or Pdgfrα lineage labels a population primarily comprised of the earliest specified precursors in the Mixl1^+^ mesendoderm niche, which can be identified in vivo and in human iPSC cell-derived directed differentiation protocol. Our findings, summarized in Figure S4, pinpoint the lineage hierarchy of specified lung precursors in gastrulating mesendoderm, further supported by the scRNA-seq analysis in early embryonic development(Pijuan-Sala et al., 2019) (Figure S4B).

As previously indicated that nascent mesoderm differentiation into a CPP fate(Bardot et al., 2017; Devine et al., 2014; Ng et al., 2022; Peng et al., 2013), we clarified the orderly mesendoderm progression of gene expression, Mixl1, Pdgfrα, and Foxa2, and lung progenitor-related markers that parallels the commitment of Foxa2^+^ Pdgfrα^+^ mesendoderm to an LPM and DE fate in the lung directed differentiation protocol using human iPSC (Figure S4). Our lineage tracing analysis also highlighted the unanticipated Foxa2 lineage program, the progressive increase of lineage labeling by spontaneous expression of *Foxa2 mRNA* that occupies more than half of the lung mesenchyme during lung development. Foxa2-lineage^+^ lung mesenchyme is a part of Pdgfrα lineage^+^ cells, potentially providing a unique competitive developmental niche during lung development. Further analyses using Foxa2CreERT2 lineage tracing mice are required to clarify it. Intriguingly, the Foxa2 lineage^+^ mesenchyme did not show any discrete fates that would predict anatomical localization—depleting Fgfr2 in the Foxa2 lineage results in the loss of the Fgfr2 mitogen-mediated function of Foxa2 lineage^+^ lung mesenchyme, which leads to the loss of proliferative ability in most host lung mesenchyme. The Foxa2 lineage labeling in lung mesenchyme is around 20% at E14.5 (Figure. 2D and 2E). Moreover, the donor cell complementation in lung mesenchyme at E14.5 still holds host-derived lung mesenchyme but decreases at E17.5. These results indicate that the defective Foxa2 lineage is critically important for efficient lung complementation.

An unambiguous proof of the developmental origins of patterned organs is critical for developing regenerative strategies and a better understanding of the genes responsible for congenital malformations. For the future development of whole lung generation via CBC using human iPSCs, the human-derived developmental program modification to match the host animals is the key. More broadly, our studies offer a new paradigm that can apply to modeling various congenital lung diseases and future autologous transplantation therapies using iPSC in the near future.

## Acknowledgments

We thank Zurab Ninish for his technical assistance. We sincerely appreciate the generous support from Dr. Hiromitsu Nakauchi at Stanford University and the considerate support and scientific input from Dr. Wellington Cardoso at the Columbia Center for Human Development (CCHD) and the members of Cardoso’s lab and CCHD. We acknowledge the support from the CCHD Medicine Microscopy core (MMC), Columbia Stem Cell Initiative (CSCI) Flow Cytometry core (SONY MA900), and Genetically Modified Mouse Model Shared Resource (GMMMSR) for blastocyst injection. This work was funded by NIH-NHLBI 1R01 HL148223-01, DoD PR190557, PR191133 to M. M., JSPS202080340, and The Uehara Memorial Foundation to A. M.

## Author contributions

A.M. and M.M. designed all experiments; Z.N. and A.M. maintained mutant mice for the injection; C.S.L. performed blastocyst injection and embryo transfer; J.T., A.S., Y.S., Y.H., H.S., supported lineage-tracing, chimera analyses, and genotyping; H.S., D.S., and S. T. helped to generate mouse and human iPSCs, J.S. and F.H. kept human iPSC-directed differentiation, N.D. provided Foxa2^Cre/+^ mice, A.M. and M.M. wrote the paper; Y.H., H.S., J.W., J.Q., and F.H. gave crucial insights on the experiments and the manuscripts. There is no competing financial interest.

## Methods

### Mouse

*Shh*^*Cre/+*^ mice (cat. 05622), *Rosa26*^*tdTomato/tdTomato*^ mice (cat. 07914), Rosa26^nT-nG/nT-nG^ mice (cat. 023035) and *Pdgfrα*^*CreERT2/+*^ mice (cat. 032770) were obtained from the Jackson Lab. X. Zhang kindly gifted *Fgfr2*^*flox/flox*^ mice. We further backcrossed these mice for over three generations with CD-1 mice (cat. 022) from the Charles River. Dr. Nicole C Dubois kindly provided *Foxa2*^*Cre/Cre*^ mice (129xB6 mixed background). For conditional deletion of *Fgfr2* (*Fgfr2*^*cnull*^), we crossed *Fgfr2*^*flox/flox*^; *Rosa26*^*tdTomato/tdTomato*^ females with *Foxa2*^*Cre/Cre*^; *Fgfr2*^*flox/+*^, *Foxa2*^*Cre/+*^; *Fgfr2*^*flox/+*^ or *Shh*^*Cre/+*^; *Fgfr2*^*flox/+*^ males, respectively. PCR performed genotyping of the *Shh-Cre, Pdgfrα-CreERT2, Rosa26-nTnG*, and *Rosa26-tdTomato* alleles according to the protocol provided by the vendor. For the CBC, genotyping of chimeric animals was confirmed by GFP-negative sorted liver cells and lung cells. For detecting the *Fgfr2* floxed allele, we performed PCR using the primer sets: FR2-F1, 5′-ATAGGAGCAACAGGCGG-3′, and FR2-F2, 5′-CAAGAGGCGACCAGTCA-3′(Mori et al., 2019). For lineage tracing with Pdgfrα^CreERT2/+^; Rosa26^tdTomato/+^ mice, 1 dose of 200 μg tamoxifen (MedChem Express, HY-13757A) per g of body weight was given via oral gavage injection. All animal experiments were approved by Columbia University Institutional Animal Care and Use Committee in accordance with US National Institutes of Health guidelines.

### Culture of mouse iPSC

We cultured iPSC in a2i/VPA/LIF medium on a feeder, as previously reported(Mori et al., 2019). These PSC cells were passaged at a split ratio of 1:10 every 2–3 d.

### Culture of human iPSCs (hiPSCs)

All iPSC lines were maintained in feeder-free conditions on laminin iMatrix-511 silk E8 (Amsbio, AMS.892021) in StemFit 04 complete Medium (Amsbio, SFB-504), supplemented with Primocin (Invivogen, ant-pm-1), and passaged with TrypLE Select (Gibco, A1285901). All human iPSC lines used were characterized for pluripotency and were found to be karyotypically normal. The BU3NGST cell line was kindly gifted by Dr. Finn Hawkins and Dr. Darrell Kotton at Boston University, Boston, MA. Dr. Jennifer Davis, the University of Washington School of Medicine, Seattle, WA, kindly gifted the Rainbow cell line. PD2 and TD1 hiPSC were generated from deidentified commercially available human peripheral blood mononuclear cell and tracheal epithelial cell lines via the manufacturing protocol of Sendai virus-mediated reprogramming (CytoTune2.0) (ThermoFisher, A16517). Every other month, all iPSC lines screened negative for mycoplasma contamination using a MycoAlert PLUS detection kit (Lonza, LT07-710).

### Differentiation of hiPSCs into lung epithelial and mesenchymal cells

The directed differentiation protocols were modified from previous protocols to maximize lung mesenchymal cell generation concomitantly with NKX2-1^+^ lung epithelium. Briefly, DE and LPM precursors were induced once seeded hiPSC-formed colonies by the Activin induction using the STEMdiff Definitive Endoderm Kit (StemCell Technologies, 05110) for 72 hours. Differentiated cells were dissociated and passaged in Laminin511-coated tissue culture plates in a complete serum-free differentiation medium (cSFDM)(Chen et al., 2017). To induce DE and LPM into the anterior foregut endoderm and mesoderm, the cSFDM was supplemented with 10μM SB431542 (MedChem Express, HY-10431) and 2μM Dorsomorphin (Tocris, 3093) for 48 hours and 10μM SB431542 and 2μM IWP2 (Tocris, 3533) for 24 hours. Cells were then cultured for 7-10 additional days in cSFDM containing 3μM CHIR99021, 10ng/ml recombinant human FGF10 (R&D Systems, 345-FG), 10ng/ml recombinant human KGF (R&D Systems, 251-KG), 10 ng/mL recombinant human BMP4 (R&D Systems, 314-BP), and 50nM retinoid acid (Sigma-Aldrich, R2625) to induce NKX2-1 positive lung epithelial cells and WNT2^+^TBX4^+^ lung mesenchymal cells.

### Immunofluorescence (IF)

Before the immunostaining, antigen retrieval was performed using Unmasking Solution (Vector Laboratories, H-3300) for 10 min at around 100 °C by microwave. 7-μm tissue sections were incubated with primary antibodies in the buffer of M.O.M. kit (Vector Laboratories, MKB-2213-1) overnight at 4 °C, washed in PBS, and incubated with secondary antibodies conjugated with Alexa488, 567, or 647 (ThermoScientific, 1:400) with NucBlue Fixed Cell Ready Probes Reagent (Hoechst) (ThermoScientific, R37605) for 1.5 h, and mounted with ProLong Gold antifade reagent (Invitrogen, P36930). The images were captured by a Zeiss confocal 710 microscopy. The antibodies were listed in the Supplemental Table. 1.

### Immunocytochemistry

Cells on culture dishes were fixed with 4% Paraformaldehyde (PFA) for 30 min at room temperature (RT), permeabilized, and blocked with staining buffer containing 0.025% Triton X-100 and 1% BSA for 1 hour at RT. Primary antibodies (Supplementary Table 1) were incubated overnight at 4 °C in the staining buffer. After three washes in PBS, secondary antibodies (Supplementary Table 1) and NucBlue Fixed Cell Ready Probes Reagent (Hoechst) were incubated for 1 h. The samples were imaged using DMi8 Leica widefield microscope. The antibodies were listed in the Supplemental Table. 1.

### RNAScope in situ hybridization (ISH)

RNA at E18.5 lung sections were stained by RNAScope probes: Mm-Foxa2-T8 (Advanced Cell Diagnostics, #409111-T8) or Negative control (NC) (Advanced Cell Diagnostics, #324341) using the RNAScope HiPlex12 Reagent kit v2 (Advanced Cell Diagnostics, #324419) according to manufacture-provided protocol. Then, sections were incubated with tdTomato antibody for 2 hours at room temperature, washed in PBS, and incubated with secondary antibody conjugated with Alexa488 with NucBlue Fixed Cell Ready Probes Reagent (Hoechst) for 1.5 h, and mounted with ProLong Gold antifade reagent. The images of *Foxa2* and NC ISH were captured with the same setting by a Zeiss confocal 710 microscopy.

### Flow cytometry (FCM) analyses of mouse lung tissue

Lungs from lineage tracing mice at E14.5, E18.5, P0, four weeks or lungs from CBC chimeric mice at E14.5 and E17.5 were harvested and prepared for the FCM, as previously described(Mori et al., 2019). Briefly, tissues were minced with microscissors, and 1 ml of pre-warmed dissociation buffer (1 mg/ml DNase (Sigma, DN25), 5 mg/ml collagen (Roche, 10103578001), and 15 U/ml Dispase II (Stemcell Technologies, 7913) in HBSS), incubated at 37 °C on the rocker with 50 r.p.m. speed, and neutralized with the dissociation buffer by FACS buffer containing 2% FBS, Glutamax, 2mM EDTA and 10mM HEPES in HBSS after the 30 min incubation. After filtrating the cells with a 40-μm filter (FALCON, 352235), cell pellets were resuspended with 1 ml of cold RBC lysis buffer (Biolegend, 420301) to lyse the remaining erythrocytes for 5 min on ice, and neutralized by 1 ml cold FACS buffer. After that, it was centrifuged them at 350 rcf, 4 °C, for 3 min to remove the lysed blood cells. For FCM analysis, one million cells were transferred in 100 μl of FACS buffer supplemented with 0.5μM Y27632 and then added 2 μl Fc Block (BD Pharmingen, 553141) per sample followed by 10 min incubation on ice. Cells were incubated with the following antibodies: CD31-APC (Biolegend, 102510, 1/50), Epcam-BV711 (BioLegend, 118233, 1/50), or Epcam-BV421 (Biolegend, 118225, 1/50), Aqua Zombie (BioLegend, 423101, 1/100), CD45-BV605 (BioLegend, 103104, 1/50) for 30 min on ice. After staining, cells were washed twice with FACS buffer before resuspending in 500 μl FACS buffer for the subsequent analyses using SONY MA900 or NovoCyte. Compensation was manually performed to minimize the tdTomato signal leakage to the GFP channel using FlowJo (ver. 10. 7. 1). 2 samples from E14.5 lineage tracing mice were removed from analysis based on inaccurate staining of live staining. Total 4 lungs were calculated from 6 embryos.

### nGFP^+^iPSC establishment and preparation for CBC donor

E14.5 lung tissues of *Rosa26*^*nTnG/nTnG*^ mice (JAX, cat. 023035, C57BL/6NJ background) were harvested in a dissociation buffer described above. The dissociated cells were seeded on a 10cm dish, and only lung fibroblast survived after 1 week in MEF medium(Mori et al., 2019). The fibroblasts were passaged using Accutase (Innovative Cell Technologies, AT104) and seeded on gelatin (Millipore-Sigma, ES006B)-coated 6-well plates with a density of 0.1 million cells per well. Upon cell attachment, Yamanaka reprogramming factors were induced to iPSCs via Sendai virus using CytoTune2.0 (ThermoFisher, A16517). To establish nGFP^+^ iPSCs, the Cre plasmid was transfected using Fugene HD transfection reagent (Promega, E2311), then sorted out GFP^+^tdTomato^−^ live cells by FACS (SONYMA900), and single clones were expanded.

For the CBC donor cell preparation, nGFP^+^iPSCs cultured in a2i/VPA/LIF (Mori et al., 2019) were trypsinized and resuspended in 4 ml cold DMEM + 10% FBS immediately and filtering the cells with a 40-μm filter. Cells were centrifuged at 350 rcf, 4 °C, for 3 min, and the supernatant was removed. After being washed with flow buffer containing 0.2% BSA, 1% Glutamax, and 1μM Y27632, the cells were resuspended in 100μl/1 million cells with flow buffer. The following antibodies were added: Epcam-BV421 (1:50), SSEA1-PE (1:50), CD31-APC (1:50), Zombie Aqua Fixable Viability Kit (1:100). Epcam^high^SSEA1^high^CD31^high^ cells were sorted by FACS (SONYMA900) and subsequently prepared for the injection.

### Blastocyst preparation and embryo transfer

Blastocysts were prepared by mating *Foxa2*^*Cre/Cre*^; *Fgfr2*^*flox/+*^, *Foxa2*^*Cre/+*^; *Fgfr2*^*flox/+*^ or *Shh*^*Cre/+*^; *Fgfr2*^*flox/+*^ males (all 129 x B6 x CD-1 background) with superovulated Fgfr2^flox/flox^; Rosa26^tdTomato/tdTomato^ females (129 x B6 x CD-1 background). Blastocysts were harvested at E3.5 after superovulation(Mori et al., 2019). 20 sorted nGFP^+^iPSCs were injected into each blastocyst. After the iPSC injection, blastocysts were cultured in an M2 medium (Cosmobio) for a few hours in a 37 °C, 5% CO2 incubator for recovery. Then, blastocysts were transferred to the uterus of the pseudopregnant foster mother.

### Real-time-quantitative RT-PCR (qRT-PCR)

Total RNA was extracted using a Direct-zol™ RNA MiniPrep Plus kit (Zymo Research, R2072), and cDNA was synthesized using Primescript™ RT Master Mix (Takara, RR036B). The cDNAs were then used as templates for qRT-PCR analysis with gene-specific primers. Reactions (10 μl) were performed Luna® Universal qPCR Master Mix (New England Biolabs, M3003X). mRNA abundance for each gene was determined relative to GAPDH mRNA using the 2^−ΔΔCt^ method. The primers were listed in the Supplemental Table. 2. Data were represented as mean ± SD of measurements. The number of animals or cells per group is provided in the legends. The undetected values in each biological experiment in Fig.3d were removed from the graphs.

### Statistical analysis

Data analysis was performed using Prism 8. Data acquired by performing biological replicas of two or three independent experiments are presented as the mean ± SD. Statistical significance was determined using a two-tailed t-test and unpaired one-way or two-way ANOVA with the Tukey post hoc test. \**P* < 0.05, \*\**P* < 0.01, \*\*\**P* < 0.001, ns: non-significant.

## Data availability

scRNA-seq data for mouse gastrulation and early organogenesis described in the manuscript have been analyzed from the deposited database at https://marionilab.cruk.cam.ac.uk/MouseGastrulation2018/. The authors declare that all data supporting the results of this study are available within the paper and the Supplementary Information.

## SUPPLEMENTAL FIGURES

**Figure S1.**
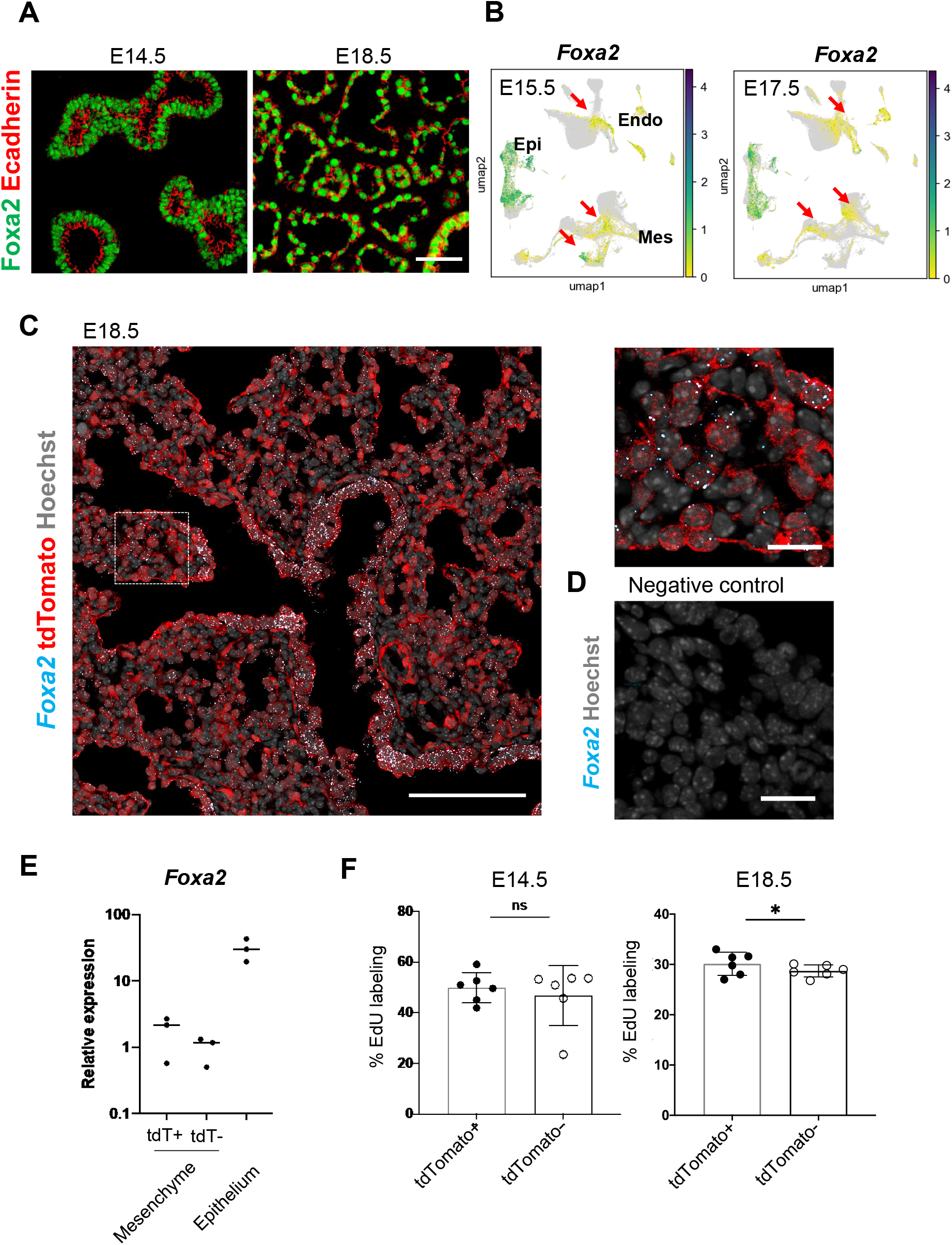
Foxa2-lineage gradually increased in the mesenchyme and endothelial cells during mouse lung development. (A) Representative IF-confocal imaging of E14.5 and E18.5 mouse embryonic lungs. Foxa2 (green) was expressed in lung epithelial cells labeled by E-cadherin (red) but not in E-cadherin negative lung mesenchymal area. Scale bar = 50μm (B) Single-cell analysis of mouse LungMAP. *Foxa2* expression was observed in lung mesenchyme and endothelium in E15.5 and E17.5 (Arrows). (C), (D) Representative IF and in situ hybridization-confocal imaging of E18.5 mouse embryonic lungs. Dotted lines: enlarged image. *Foxa2* mRNA was expressed in lung epithelial cells and lung mesenchyme. Most of the *Foxa2* mRNA expression was colocalized with tdTomato. Scale bars (left, right) = 50 μm, 20μm (E) Graphs: *Foxa2* qPCR for tdTomato (tdT)^+^ or tdTomato^−^ CD45^−^ CD31^−^ EPCAM^−^ mesenchymal cells and CD45^−^CD31^−^EpCAM^+^ epithelial cells sorted by FACS from E18.5 Foxa2-lineage tracing mice. (n = 3, each biological replicate) (F) Flow cytometry quantitative analyses: EdU labeling cells in CD31^+^Epcam^−^ endothelium at E14.5 and E18.5 (n = 6 each independent biological replicates, respectively) of *Foxa2*^*Cre/+*^; *Rosa*^*tdTomato/+*^ lungs. Statistical analysis: paired Student’s t-test, significance at \**P* < 0.05, ns: non-significant.

**Figure S2.**
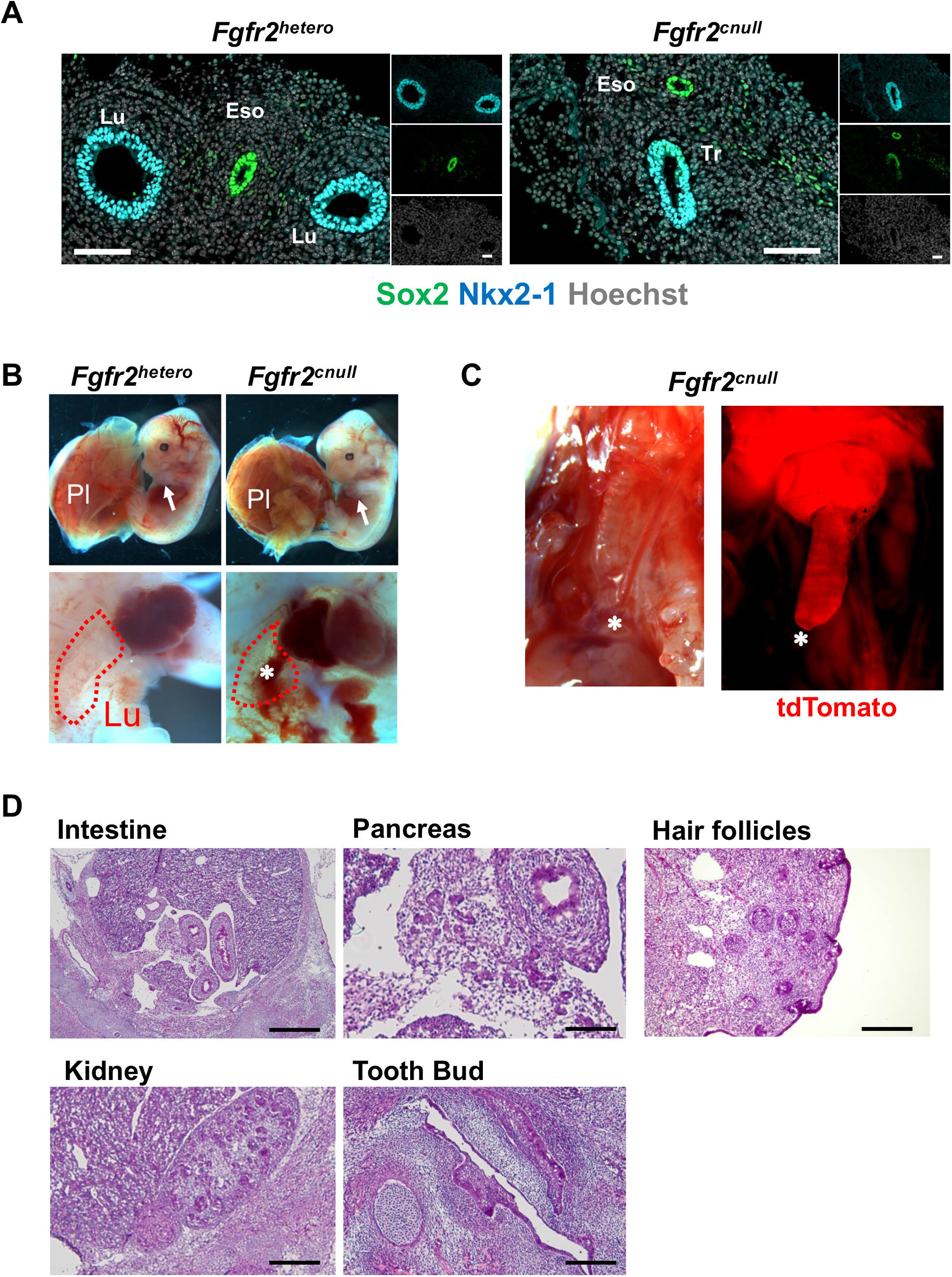
Lung agenesis phenotype in the *Foxa2*^*Cre/+*^*Fgfr2*^*cnull*^ mice. (A) Representative IF staining of E10.5 *Foxa2*^*Cre/+*^;*Fgfr2*^*hetero*^ and *Foxa2*^*Cre/+*^;*Fgfr2*^*cnull*^: *Fgfr2*^*cnull*^ did not form Sox2^+^NKX2-1^+^proximal airway bifurcation. Scale bar = 200μm. Lu: lung, Eso: Esophagus. Tr: trachea. Nkx2-1 (blue), Sox2 (green). (B) Representative Gross morphology of E14.5 *Foxa2*^*Cre/+*^;*Fgfr2*^*hetero*^ and *Foxa2*^*Cre/+*^;*Fgfr2*^*cnull*^: No difference in the appearance of the embryo between *Fgfr2*^*hetero*^ and *Fgfr2*^*cnull*^ (Top). Limb (arrows) and placenta (pl) are present in mice of all genotypes. Conversely, the *Fgfr2*^*cnull*^ embryo showed lung agenesis phenotype (Bottom, asterisk) (n = 6 per each group). (C) Representative image of *Fgfr2*^*cnull*^ lungs. Foxa2 lineage^+^ tdTomato signal indicates the trachea ends in the middle of the thoracic cavity (asterisk). (D) HE staining: The internal organs such as the intestine, pancreas, kidney, tooth buds, and hair follicles were normally formed in *Fgfr2*^*cnull*^ E16.5 embryos. Scale bar = 500μm.

**Figure S3.**
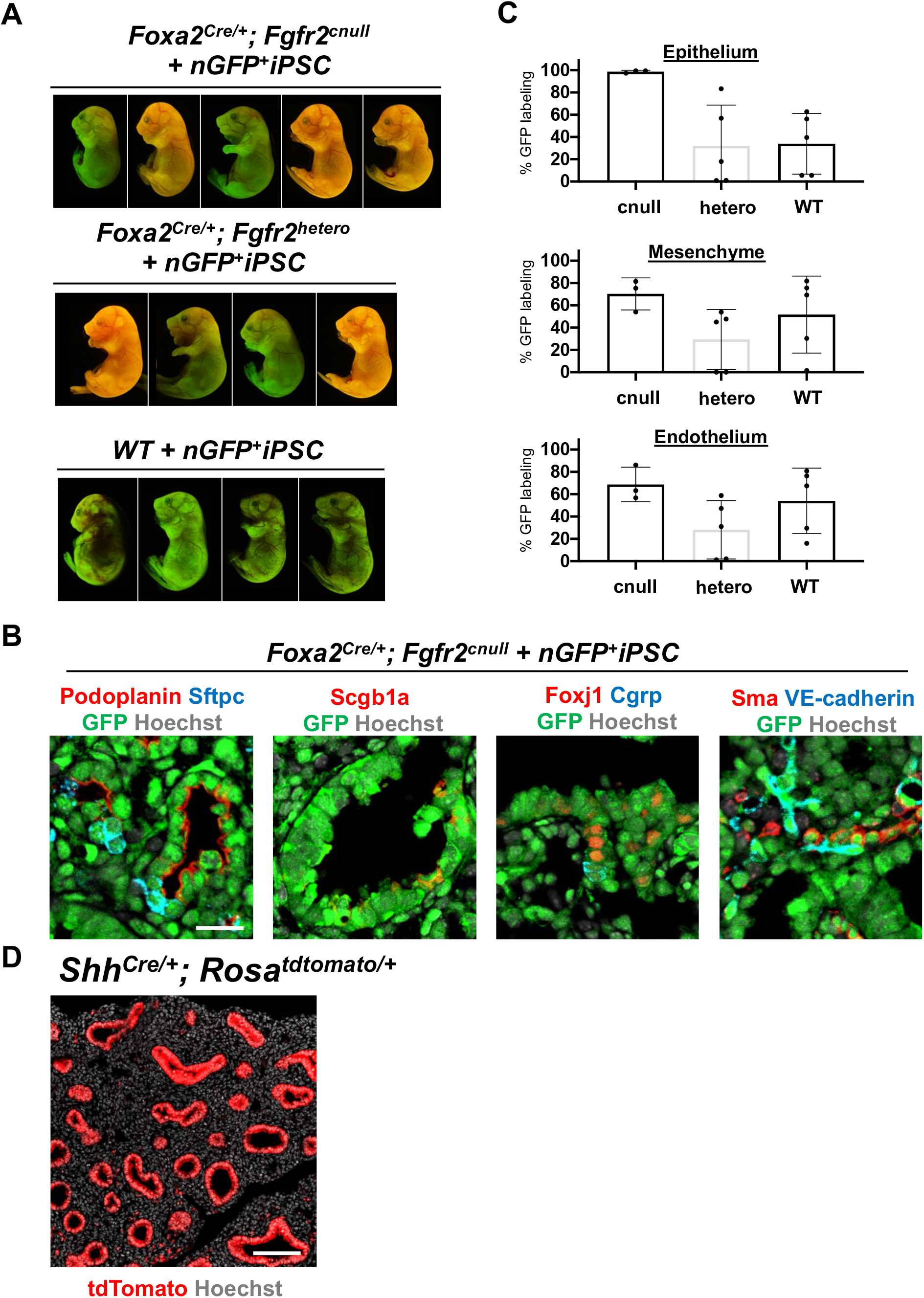
The complemented embryos of *Foxa2*^*Cre/+*^; *Fgfr2*^*cnull*^ +nGFP^+^iPSCs showed normal gross morphology and various chimerism. (A) Representative gross morphology of chimeric embryos of *WT* + nGFP^+^iPSCs, *Foxa2*^*Cre/+*^; *Fgfr2*^*hetero*^ + nGFP^+^iPSCs, *Foxa2*^*Cre/+*^; *Fgfr2*^*cnull*^ +nGFP^+^iPSCs. The color indicated various GFP^+^ chimerism on the skin of those embryos and the residual host Foxa2-lineage^+^ tdTomato signals. (B) Representative IF-confocal imaging of E17.5 lungs of chimeric *Fgfr2*^*cnull*^+ nGFP^+^iPSCs. The E17.5 rescued lungs from *Fgfr2*^*cnull*^ +nGFP^+^iPSCs expressed differentiated cell markers such as Podoplanin (alveolar type1 cells), Sftpc (type 2 cells), Scg1a1 (club cells), Foxj1 (multiciliated cells), Cgrp (neuroendocrine cells), Sma (smooth muscle cells), and VE-cadherin (endothelial cells). Scale bar = 20μm. (C) Graphs: % of donor cell chimerism: % of GFP in CD31^−^Epcam^+^ lung epithelium fraction, CD31^−^ Epcam^−^ mesenchyme, and CD31^+^Epcam^−^ endothelium analyzed by flow cytometry. Each plot: a different biological animal (see table 2). *Foxa2*^*Cre/+*^; *Fgfr2*^*cnull*^; *Rosa*^*tdTomato*^ (*n=3*, independent biological replicates), *Foxa2*^*Cre/+*^; *Fgfr2*^*hetero*^; *Rosa*^*tdTomato/+*^(*n=5*), and *WT* (*n=5*) at E14.5. (D) Representative IF-confocal imaging of E14.5 Shh-lineage tracing mouse lungs (*Shh*^*Cre/+*^; *Rosa*^*tdTomato/+*^). Shh-lineage labeled the entire lung epithelium but rarely mesenchyme, distinct from Foxa2 lineage mice (see Fig. 2F). Scale bar = 100μm.

**Figure S4.**
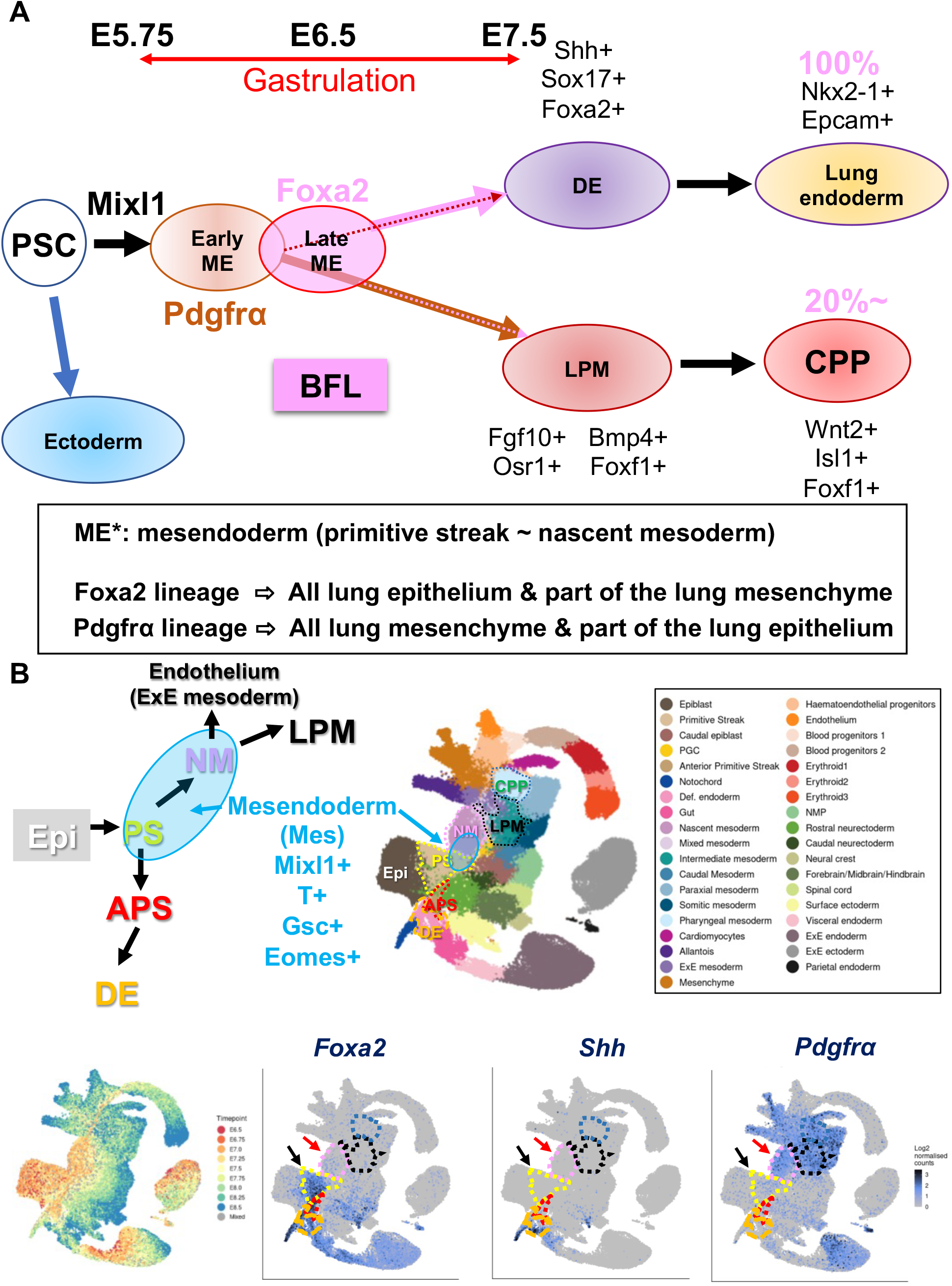
Summary of the results and proposed models. (A) Pluripotent stem cell (PSC) differentiation will be initiated by the Mixl1^+^ induction and the subsequent temporal expression of an early mesendoderm (ME) marker, Pdgfrα. After that, the late mesendoderm marker marked by Foxa2 will turn on. Pdgfrα and Foxa2-lineage partially overlap at the primitive streak stage during gastrulation. Pdgfrα^+^ early ME lineage gives rise to the partial lung epithelial cells, most likely by the overlapping Foxa2^+^ lineage and entire lung mesenchyme through lateral plate mesoderm (LPM) induction and cardiopulmonary lineage (CPP). On the other hand, the Foxa2-lineage gives rise to the whole lung epithelium and about 20% of CPP in early lung development at E12.5∼E14.5. Strikingly, the whole lungs, including lung epithelium, mesenchyme, and endothelial cells, were produced solely by donor iPSCs via complementing the Foxa2-lineage’s mitotic defective lung niches. It proves that a Foxa2-lineage works as a bona fide lung generative lineage (BFL), enough to generate entire lungs by donor iPSCs via CBC. (B) Top left panel: Schematic diagram of mesendoderm lineage trajectory for lung formation based on the single cell RNA-seq (scRNA-seq) deposited database(Pijuan-Sala et al., 2019). During gastrulation, a bipotent mesendoderm (Mes: rounded blue area) appears in the transition from PS and NM, labeled by *Mixl1, T, Gsc, and Eomes*. NM forms lateral plate mesoderm (LPM). Mesendoderm also gives rise to anteriolar PS (APS) and, subsequently, definitive endoderm (DE). Top right panel: Clustering of scRNA-seq provided in the deposited database(Pijuan-Sala et al., 2019). Bottom left panel: Timepoint of scRNA-seq provided in the deposited database(Pijuan-Sala et al., 2019). bottom: *Foxa2* is expressed in the PS (black arrow) and the part of NM (red arrow). In contrast, *Shh* appeared in DE but not in PS, NM, or APS. A few cells showed *Pdgfrα* in PS (black arrow) but most expressed *Pdgfrα* in NM (red arrow) and LPM.

